# Prediction of protein assemblies by structure sampling followed by interface-focused scoring

**DOI:** 10.1101/2023.03.07.531468

**Authors:** Kliment Olechnovič, Lukas Valančauskas, Justas Dapkūnas, Česlovas Venclovas

**Affiliations:** Institute of Biotechnology, Life Sciences Center, Vilnius University, Saulėtekio 7, LT-10257 Vilnius, Lithuania

**Keywords:** Protein-protein interactions, protein complexes, AlphaFold-multimer, docking, estimation of model accuracy, interface scoring, VoroIF-jury

## Abstract

Proteins often function as part of permanent or transient multimeric complexes, and understanding function of these assemblies requires knowledge of their three-dimensional structures. While the ability of AlphaFold to predict structures of individual proteins with unprecedented accuracy has revolutionized structural biology, modeling structures of protein assemblies remains challenging. To address this challenge, we developed a protocol for predicting structures of protein complexes involving model sampling followed by scoring focused on the subunit-subunit interaction interface. In this protocol, we diversified AlphaFold models by varying construction and pairing of multiple sequence alignments as well as increasing the number of recycles. In cases when AlphaFold failed to assemble a full protein complex or produced unreliable results, additional diverse models were constructed by docking of monomers or subcomplexes. All the models were then scored using a newly developed method, VoroIF-jury, which relies only on structural information. Notably, VoroIF-jury is independent of AlphaFold self-assessment scores and therefore can be used to rank models originating from different structure prediction methods. We tested our protocol in CASP15 and obtained top results, significantly outperforming the standard AlphaFold-Multimer pipeline. Analysis of our results showed that the accuracy of our assembly models was capped mainly by structure sampling rather than model scoring. This observation suggests that better sampling, especially for the antibody-antigen complexes, may lead to further improvement. Our protocol is expected to be useful for modeling and/or scoring protein assemblies.

## Introduction

The recently developed AlphaFold method has revolutionized protein structure prediction, achieving unprecedented accuracy in modeling individual proteins^1, 2^. Application of AlphaFold enabled construction of structural models for nearly all currently known proteins^3, 4^. However, many proteins function not as single chains, but as components of multichain protein assemblies (complexes). The prediction of structures for protein assemblies remains a significant challenge, because it requires accurately predicting not only the structures of individual subunits, but also their mutual arrangement. Although the initial release of AlphaFold was not designed to model structures of protein assemblies, it was soon discovered that the method could be ‘hacked’ to perform this task^5^. Subsequently, DeepMind released AlphaFold-Multimer, an updated AlphaFold version, specifically retrained for predicting protein complexes^6^. The authors showed that AlphaFold-Multimer outperforms the original AlphaFold in providing high accuracy predictions for protein complexes. Despite significant advance brought about by AlphaFold-Multimer, it became clear that there is still a lot of room for improvement. For example, an accurate prediction of antibody-antigen binding appeared to be problematic^6, 7^. In addition, large multisubunit complexes often turned out to be difficult modeling targets^8^. The accuracy of AlphaFold-Multimer decreases with the number of chains, and the available GPU memory limits the size of protein complexes that can be modelled. In such cases, the prediction of subcomplexes followed by their assembly into a larger complex was shown to be a viable approach^8^.

These observations prompted us to develop a protocol for the structure prediction of protein complexes, utilizing the strengths of AlphaFold at the same time addressing its known weaknesses. We reasoned that in cases, when AlphaFold-Multimer fails to generate a reliable structure of the entire protein complex, the latter could be assembled by docking AlphaFold-generated monomers or subcomplexes. As docking components could be expected to have high accuracy, the major focus should then be on the identification of docking solutions having native protein-protein interfaces.

Our protocol is based on generating a diverse set of models for a given protein assembly followed by interface-focused model scoring. Structural models of protein complexes were typically generated using AlphaFold/AlphaFold-Multimer alone or combined with docking. Complexes were then selected using the VoroIF-jury algorithm, which identifies models favored by several accuracy estimation methods developed in our group and is independent of AlphaFold-Multimer confidence scores.

Here, we describe our protocol, provide an overview of the results obtained during 15^th^ community-wide experiment on protein structure prediction (CASP15) and discuss what worked well and what needs to be improved.

## Methods

### Modeling outline

The general workflow for 3D structure prediction is illustrated in Figure 1. It consisted of two major stages: (1) construction of an ensemble of multiple diverse structural models and (2) selection of the best models using a newly developed accuracy estimation protocol.

**Figure 1.**
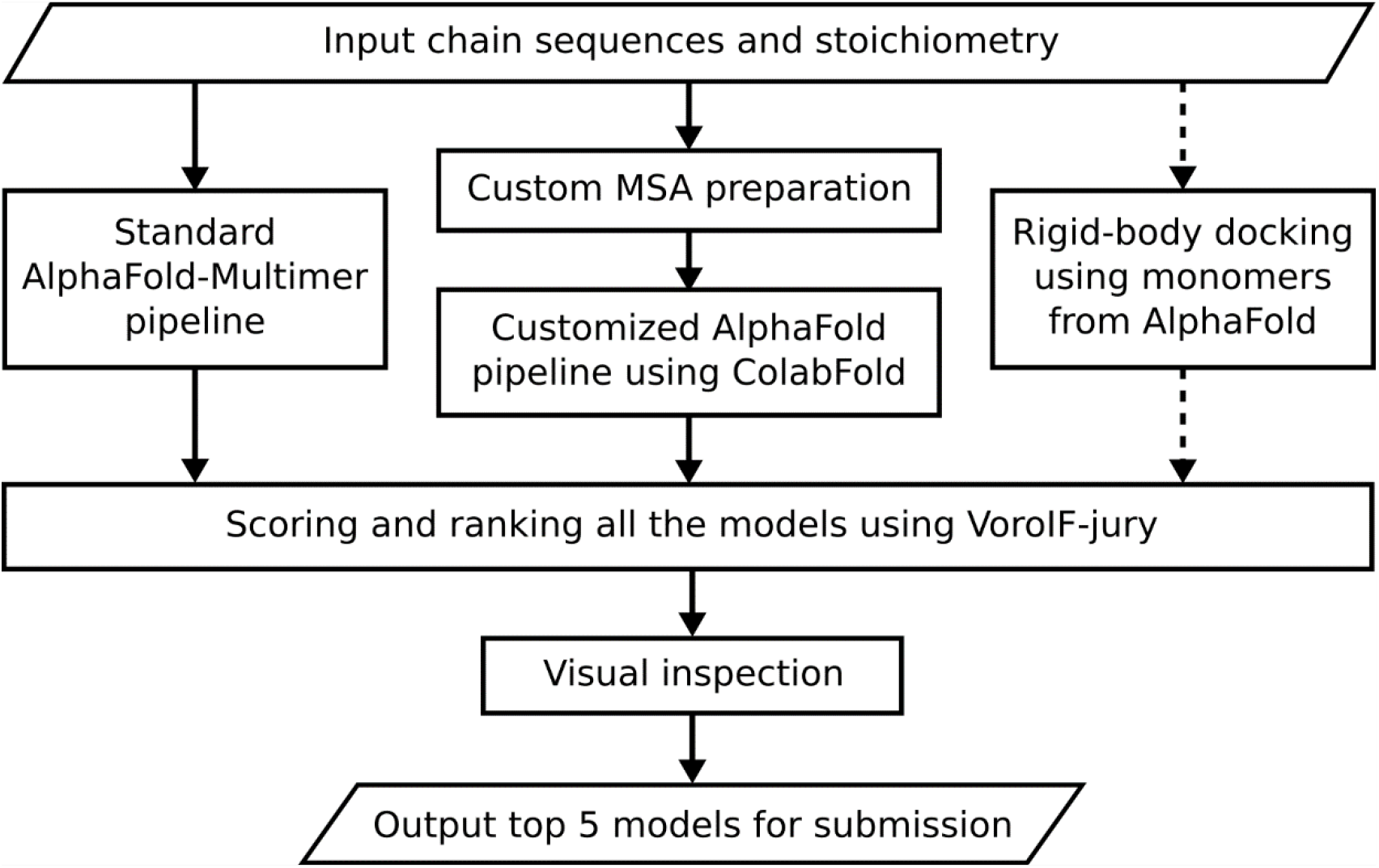
The workflow of our protein assembly modeling protocol in CASP15

For each target the initial models were generated using the standard AlphaFold from DeepMind^2, 6^, followed by customized AlphaFold pipeline using the ColabFold implementation^5^. The available structural data (homologous protein structures, disorder predictions, data published in scientific literature) were also queried for all target proteins. If AlphaFold models had high self-estimated accuracy, showed strong structural consensus and were consistent with the data for homologous proteins, no further modeling was done.

In cases when AlphaFold failed to generate the full complex (assembly was too large to handle or subunits did not form a complex), structural models were obtained using docking. Docking models were also added if the resulting AlphaFold models had poor self-estimated accuracy (pLDDT, pTM, ipTM) or were structurally diverse.

Manual interventions were used for very large targets, for which full structure models could not be produced using AlphaFold, and for several smaller targets, for which we wanted to combine information derived by several approaches. Usually this was a combination of AlphaFold modeling and docking, AlphaFold and homology modeling or diversification of the models by manually rewiring protein chains in the structures.

### Modeling of protein complexes using AlphaFold

Initial ensembles of protein complexes were constructed using AlphaFold^2^ available either as the original DeepMind’s or the ColabFold^5^ implementation. The DeepMind’s AlphaFold- Multimer v2^6^ modeling pipeline included full and reduced database presets for multiple sequence alignment construction and PDB templates and was used to generate the initial models for each target.

The ColabFold-based AlphaFold modeling pipeline employed a variety of different parameters and conditions to achieve extensive structure sampling. More specifically, each assembly was modeled using a number of custom multiple sequence alignments (MSAs) consisting of different sequence sources and pairing methods. These multiple MSAs were used as an input for both AlphaFold-Multimer v2 and AlphaFold-pTM neural network models with “residue index offset trick”. Overall, for each of the targets, 6 different MSAs were generated and included two sequence sources, ColabFoldDB^5^ and ColabFoldDB supplemented with results from additional databases, as well as all 3 pairing methods (see Fig. 2 and text below).

**Figure 2.**
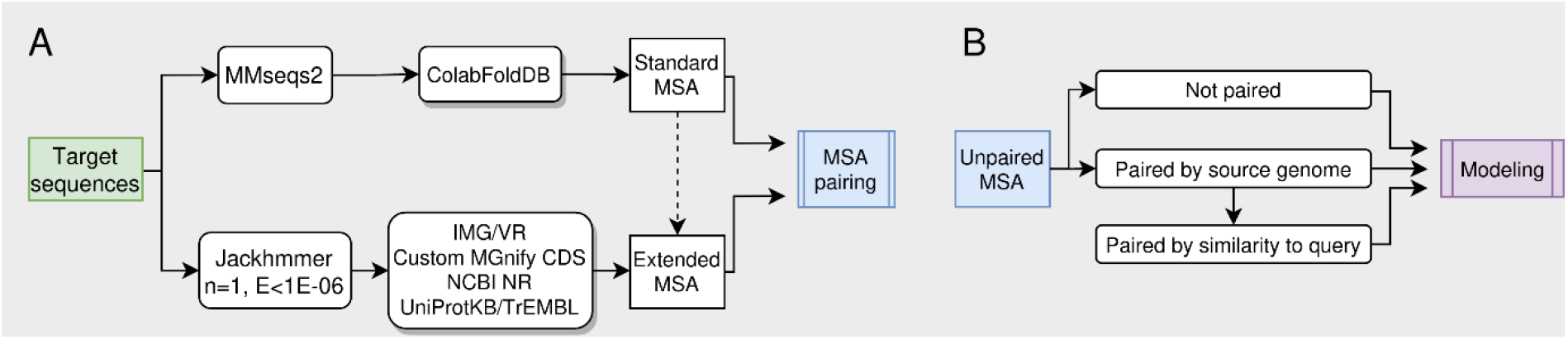
Schematic workflow for (A) generating custom multiple sequence alignments and (B) their pairing.

Additional sampling of models resulted from varying number of AlphaFold recycles. The exact number of recycles depended on the total number of residues in the assembly. Larger assemblies were usually subjected to smaller number of recycles to reduce computational costs. Typically, 3-15 recycles were applied.

### Custom multiple sequence alignment preparation

Two custom multiple sequence alignments were prepared for ColabFold (Fig. 2A). Baseline MSA was generated as in standard ColabFold using sequence search with MMseqs2 in the ColabFoldDB. Baseline MSA was then supplemented by jackhammer (HMMER 3.3.2)^9^ search over additional sequence databases including IMG/VR v3^10^, custom Mgnify redundant predicted CDS DB containing protein sequences from assemblies and genomes DB (updated on 2022-02-01)^11^, UniProt-TrEMBL (release 2021_04)^12^ and NR (2021-11-21)^13^. Jackhmmer search parameters were adopted from the original AlphaFold implementation (number of iterations = 1, inclusion E-value = 1E-06, other parameters left as HMMER defaults). In the exceptional cases, when the number of collected sequences was deemed too low, the Jackhmmer inclusion E-value was increased from default 1E-06 to 1E-03.

### Custom pairing of multiple sequence alignments

Three MSA pairing methods were implemented (Fig. 2B): (1) not paired (“checkerboard MSA”), (2) genome association pairing, and (3) genome association followed by similarity pairing. Genome association pairing consisted of pairing sequences if they came from the same genome, assembly or contig. This sort of pairing was only implemented for sequences gathered using jackhmmer from IMG/VR, custom Mgnify CDS DB, UniProt-TrEMBL and NR databases. Genome association and similarity pairing consisted of using genome paired MSA and then pairing remaining unpaired sequences according to their similarity to respective query sequences, i.e., the most similar sequence from one chain was paired with the most similar one from another chain.

### Docking

Docking was performed using established rigid-body docking tools. FTDOCK^14^ and HEX^15^ were used for generating heterodimers, whereas SAM^16^ was used for generating symmetric homomers. We did not use any internal scoring of the employed docking tools either to filter out or rank generated complexes. Docking of two components usually resulted in about 90 000 multimeric models from FTDOCK and about 6000 from Hex. Symmetric docking with SAM usually produced several hundreds of models for a single input monomer.

### Model ranking and selection

Model ranking and selection was done using the newly developed VoroIF-jury (Voronoi-based InterFace jury) procedure (Fig. 3). VoroIF-jury resembles EMA-jury, developed previously for assessing the models in recent CASP-commons experiment focused on SARS-CoV-2^17^. Currently, VoroIF-jury uses rankings from several methods, all based only on structural information, however, it can be easily extended to incorporate sequence-based or any other rankings. Given a set of models, VoroIF-jury (a) computes multiple rankings using different interface-focused scores (most of them based on the VoroMQA interface energy^18^); (b) pools the top 1, top 2, …., top N models selected by each EMA ranking into N corresponding supersets; (c) for every model in each superset calculates the VoroIF-jury interface consensus score, which is an average of the interface CAD-score^19^ values derived by comparing a given model with other models in the superset; (d) ranks models by the best achieved VoroIF-jury score; (e) removes redundant models from the final ranking using the interface CAD-score-based clustering (Fig. 3, see Supplementary Methods for algorithm details). When utilizing docking, VoroIF-jury was first applied to select top 300 from all the docking models, often exceeding 100 000. After relaxing those 300 models using OpenMM^20^ to remove clashes, the docking models were pooled together with the available AlphaFold models, and VoroIF-jury was run once again to select the final top five models.

**Figure 3.**
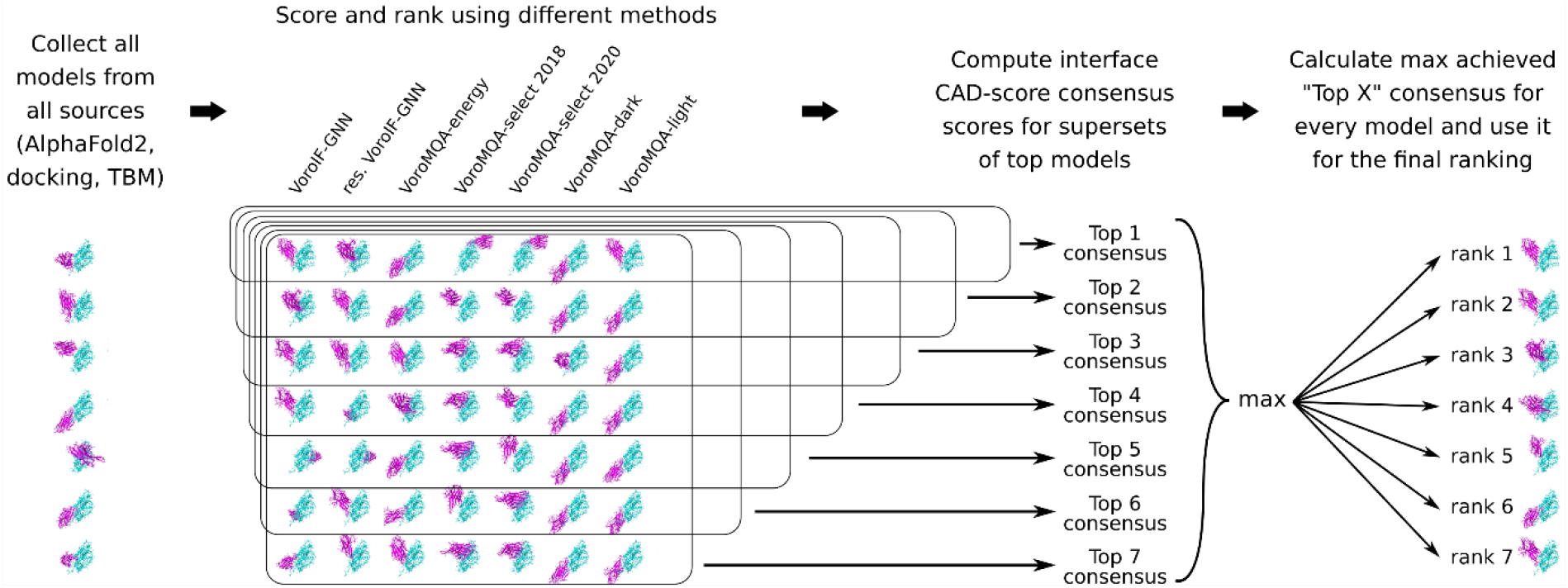
An overall scheme of the VoroIF-jury method for ranking models of protein complexes

VoroIF-jury included two newly developed interface scoring methods. The first one, a generic interatomic contact area-based energy potential, applicable for scoring of not only protein-protein, but also protein-nucleic acids interfaces, was derived from the protein-protein VoroMQA potential^18^. The second one, VoroIF-GNN, a graph neural network-based method, was developed for predicting the residue-level interface accuracy in models of protein-protein complexes (manuscript in preparation). VoroIF-GNN is based on a graph attention network (GAT) that accepts a Voronoi tessellation-derived graph of inter-chain interface contacts. The network was trained using heterodimeric models produced by rigid body redocking of complexes from PDB. The ground truth interface quality scores were calculated by comparing the models with the corresponding experimental structures using the interface CAD-score^19^. In CASP15, VoroIF-GNN was also registered as a model accuracy estimation group under the name “VoroIF”, and was among the top single-model methods in terms of the model selection performance (see the EMA assessment paper in this issue).

### Analysis of the available structural data and generation of template-based models

In addition to automatic modeling, available information on the target proteins was also considered, starting with the UniProt database^12^. Structural data for homologous proteins were queried using HHpred^21^, COMER^22, 23^ and DALI^24^ servers. Multimeric templates were identified using the PPI3D server^25^, and the oligomeric states of structurally resolved homologs were additionally checked using the RCSB PDB Advanced search^26^. If multimeric structural templates were available, homology models were generated using the PPI3D web server^27, 28^. Disordered regions were predicted by DISOPRED3^29^. All the available information was used in manual model selection and re-ranking for harder targets.

## Results

### Overview of the results

To analyze our performance in CASP15, we employed four reference-based scores also used by the CASP assessors: Interface contact similarity score (ICS or F1-score)^30^, Interface patch similarity score (IPS or Jaccard coefficient)^30^, lDDT-oligo^31^, and TM-score^32^. ICS and IPS measure the intersubunit interface accuracy, whereas lDDT-oligo and TM-score mostly reflect the overall structural accuracy. The results regarding both interface and global accuracy of our most confident CASP15 models (labeled as ‘model 1’) are summarized in Figure 4 (detailed information is provided in Supplementary Table S1). For comparison, we also included corresponding data of our CASP14 models^28^.

**Figure 4.**
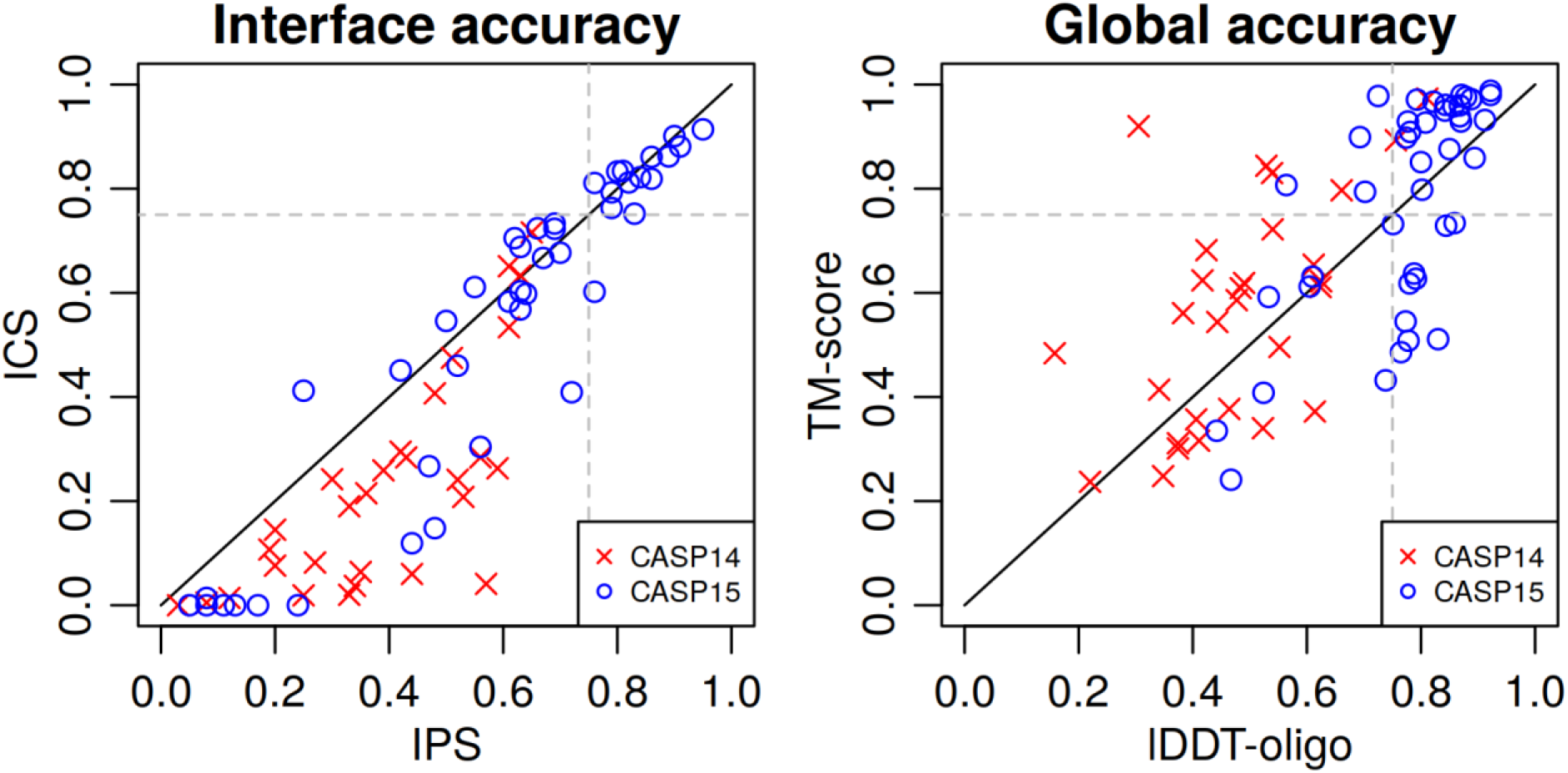
Modeling results of our group (“Venclovas”) representing the interface accuracy (left) and the overall structure accuracy (right) in CASP15 and CASP14. Broken lines mark the 0.75 threshold for all scores.

CASP15 results show a significant improvement compared to CASP14. A significant fraction of CASP15 models (14 of 43, 33%) have both ICS and IPS values over 0.75, indicating high accuracy of interface prediction. In contrast, none of the CASP14 models pass this accuracy threshold. Furthermore, in CASP15, there is a visible shift from prediction of approximate binding site (measured by IPS) to more accurate prediction of residue-residue contacts across the interface (ICS). A similar improvement can be seen in global model accuracy. A large fraction of CASP15 models (22 of 43, 51%) has both lDDT and TM-score values above 0.75, in comparison to only a couple of CASP14 models. Undoubtedly, the ability of AlphaFold to accurately predict both intrachain and interchain contacts was the main source of improvement.

Fig. 4 also shows that for seven CASP15 target assemblies, the interface contact prediction failed completely (ICS ∼ 0). Four of those cases represent either antigen-antibody or protein-nanobody complexes, a known class of complexes, for which AlphaFold-Multimer was found to perform relatively poorly^6, 7^. Two other failed cases represent structure swaps (T1176 and T1176v1). More detailed analysis of all these failed cases is presented in a separate section below.

To summarize, our CASP15 results show a significant advancement compared to CASP14. However, there is still room for improvement, in particular for certain classes of protein assemblies.

### Modeling results in the context of other CASP15 groups

Next, we compared the performance of our group with that of several other CASP15 groups. Again, we considered the models designated as first (model 1) and converted their raw reference-based scores into non-negative z-scores using the algorithm employed by the CASP organizers. We then summed the z-scores of our group (“Venclovas”), two other top-performing groups (“Zheng” and “Wallner”), and the baseline group that was running the standard AlphaFold-Multimer (“NBIS-AF2-multimer”). We also included our automated model selection protocols (the older “VoroMQA-select-2020” and the latest “VoroIF-jury”) that were registered as individual groups in the estimation of model accuracy (EMA) category. These virtual EMA groups were allowed to make selection from all the models submitted by all the CASP15 predictors. Figure 5 shows the sums of z-scores for three sets of targets: all 43 targets, 14 ‘Easy’ targets, and 29 targets classified as ‘Medium’ and ‘Hard’. Analogous plots for the sums of raw scores are presented in Supplementary Figure S1. For a compact illustration of per-target performance we plotted cumulative sums (for both z-scores and raw scores) in Supplementary Figure S2.

**Figure 5.**
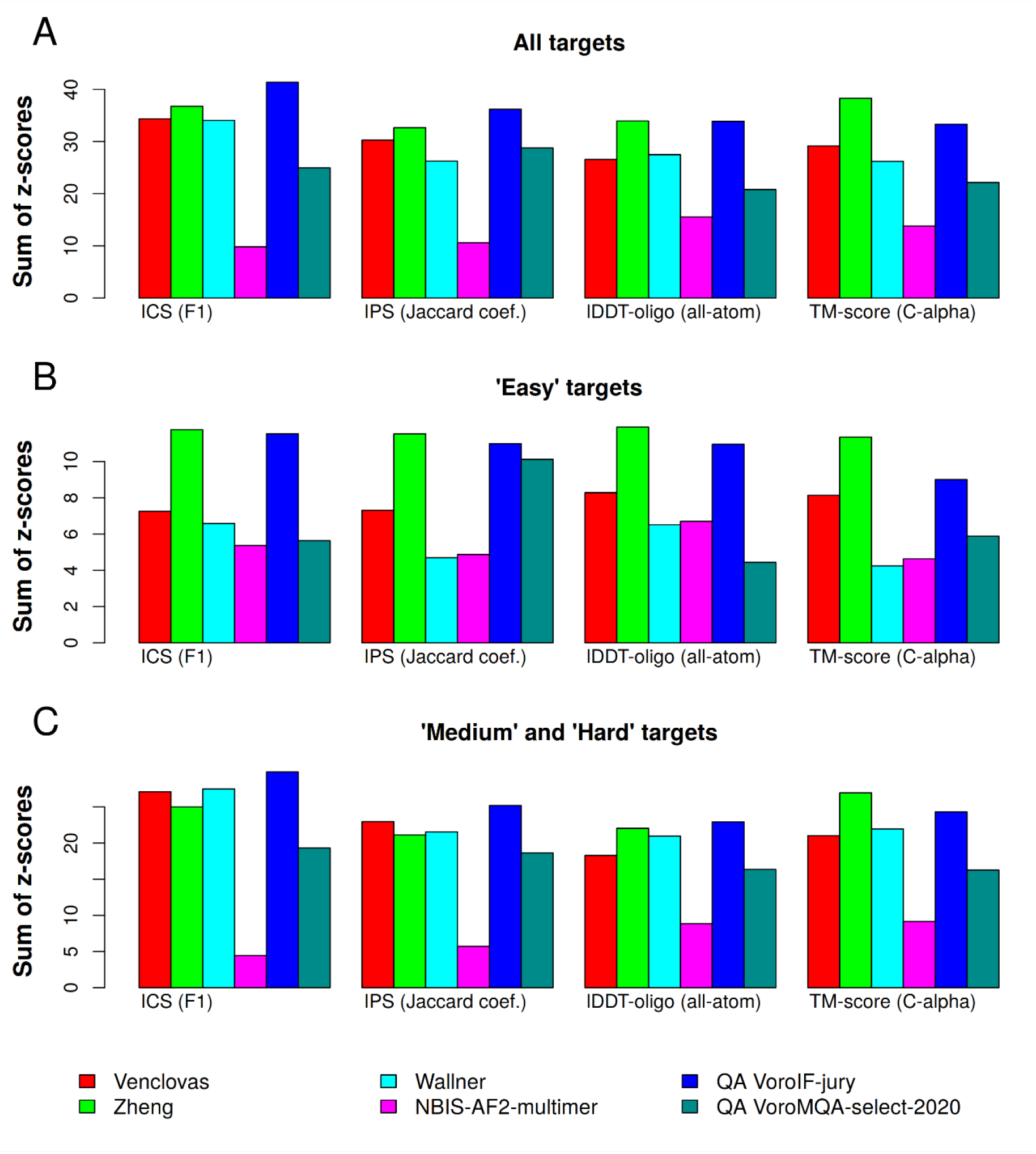
Comparison of results of our group (“Venclovas”) with the other two top-performing CASP15 groups (“Zheng” and “Wallner”) and with the baseline group that was running the standard AlphaFold-Multimer (“NBIS-AF2-multimer”). Our two automated model selection protocols (“VoroIF-jury” and “VoroMQA-select-2020”) are also included in the comparison. Performance is compared on (A) all 43 targets (B) 14 ‘Easy’ targets, and (C) 29 ‘Medium’ and ‘Hard’ targets.

Comparison shows that our group and two other top CASP15 groups clearly outperformed the standard version of AlphaFold-Multimer. The advantage is especially evident for ‘Medium’ and ‘Hard’ targets. Among the top three assembly groups, “Zheng” has the highest sums of either z-scores or raw scores when considering all the targets. This is mostly due to ‘Easy’ targets, because for ‘Medium’ and ‘Hard’ targets our group performed slightly better in terms of interface-focused scores (ICS and IPS).

### The significance of sampling and scoring

An important question regarding our results is whether the structural sampling was sufficient and whether the model scoring was effective. As can be seen in Figure 5, our automatic model scoring method, “VoroIF-jury”, when allowed to select from all CASP15 models, outperformed all the top groups based on the interface accuracy on all as well as on ‘Medium’ and ‘Hard’ targets. This clearly suggests that had there been models of higher accuracy present in our model sets, “VoroIF-jury” would have identified them. In a similar comparison based on CASP14 results^28^, our automatic selection method included here as “VoroMQA-select-2020” was inferior to the top two ‘human’ predictor groups including our own. This means that the scoring methodology of our group improved significantly since CASP14 and that the most promising route to further improvement is through better sampling.

To further investigate the impact of sampling and scoring, we analyzed our failures in predicting the correct interface. Specifically, we looked into nine targets, for which ICS score of our models was below 0.2 (Figure 4; Supplementary Table S2). We wanted to see if we had any better models in our full model set (beyond the top five models) that were missed during selection. To this end, we used the interface CAD-score^19^, an analog of ICS, because ICS scores were not available for those our models that were not submitted to CASP. We observed that for 8 of 9 targets there were no accurate models in our datasets (Supplementary Table S2). In other words, except for one case, in which model selection failed, we had nothing to choose from. Again, this indicates insufficient sampling during the modeling stage of our protocol. Interestingly, the best models for these targets came from different CASP15 groups suggesting that different modeling methods might complement each other in structure sampling.

### AlphaFold produces better models than template-based modeling

Previously, the classical template-based (homology) modeling using the available multimeric structural templates was the best possible method to predict the structures of protein complexes^28, 33–35^. To compare it with AlphaFold-Multimer, when structural templates were available, we were including a homology model generated as in CASP14^28^ among our top 5 CASP15 predictions. We compared these template-based models to our AlphaFold models, and found that AlphaFold outperforms the multimeric template-based modeling by a large margin (Table S3, Figure S3). The improvements are both at the global level and at the interaction interfaces, as indicated by higher TM-score and ICS values of AlphaFold models. Moreover, AlphaFold-Multimer generated accurate models even when homologous protein complexes were not available in the PDB (Fig. S4).

Thus, the template-based modeling can nowadays serve as a means to evaluate the available structural data for the proteins of interest, but not as the main method to obtain structure models. For example, an automated scoring combined with the analysis of homologous protein complexes facilitated selection of models for targets T1109 and T1110 (Fig. S5). T1109 is a mutant isocyanide hydratase from *Ralstonia solanacearum*, which forms a dimer different from that of the wild-type protein (T1110). We identified two possible dimers among the homologous protein structures using the PPI3D web server. Interestingly, the top scoring T1110 models corresponded to only one of these dimers, while both conformations were observed among the top models for T1109. Therefore, we hypothesized that T1109 should have an alternative dimer structure, and manually re-ranked this interface as the first model. This led to high accuracy models for both of these targets.

### Modeling of large protein assemblies

AlphaFold-Multimer failed to produce complete structures for the large multimeric targets (H1111, H1114, T1115, H1135, H1137). Nevertheless, we successfully assembled models for these challenging targets from partial AlphaFold models or by combining AlphaFold with docking (Fig. 6). For example, T1115, human stomatin, is a large ring containing 16 subunits (Fig. 6A). AlphaFold-Multimer could model only part of this ring comprised of the C-terminal domain of the protein, therefore the remaining part of the structure was modeled using symmetry docking, resulting in a fairly accurate model (TM-score=0.90 and ICS=0.68). Similarly, a combination of AlphaFold modeling and docking allowed us to obtain a model of medium accuracy for H1135 (TM-score=0.51, ICS=0.60).

**Figure 6.**
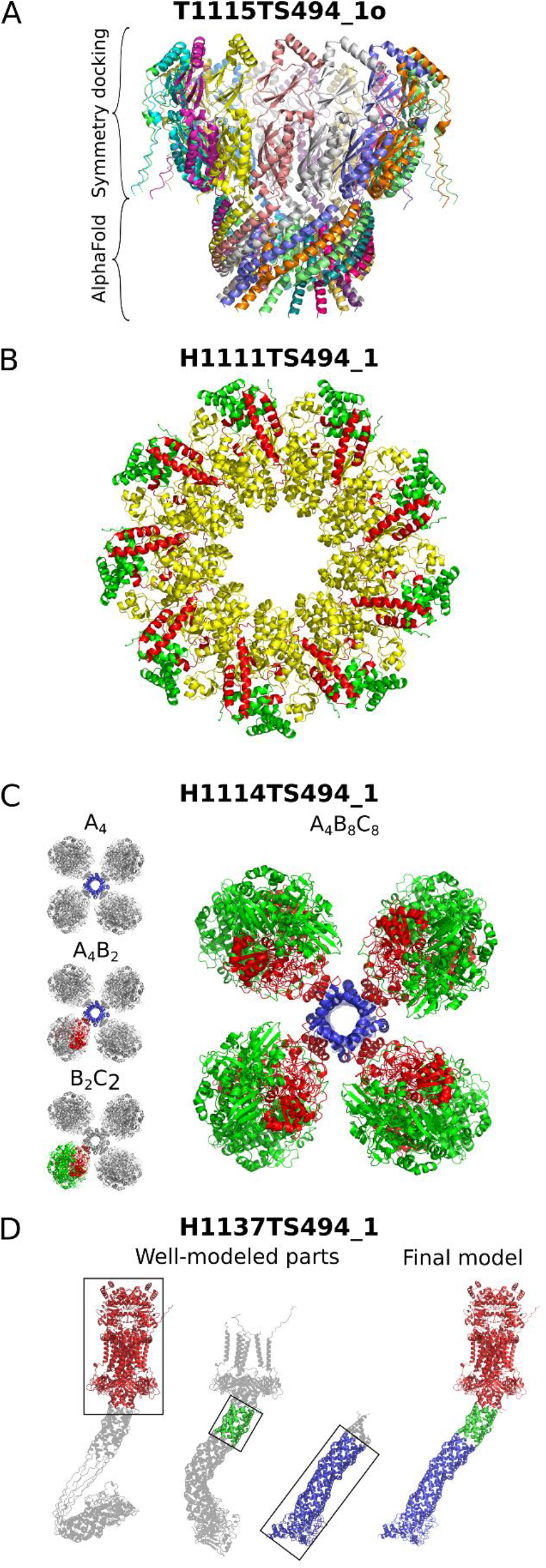
Successful assembly of large protein complexes using partial AlphaFold models. A, Homo-16-meric model for T1115 obtained by combining AlphaFold and symmetry docking; B, Model for H1111 (A_9_B_9_C_9_), obtained by aligning ABC_2_ subcomplexes (red, green and yellow) to a known C_9_ structure (yellow ring); C, Model for H1114 (A_4_B_8_C_8_; blue, red and green, respectively) combined from partial models for A_4_, A_4_B_2_ and B_2_C_2_; D, H1137 model combined from partial multimeric models.

For the remaining three large targets we modeled partial structures using AlphaFold and then combined them to obtain the full protein complex. In the case of H1111, a 27-mer having the A_9_B_9_C_9_ stoichiometry, AlphaFold models for ABC_2_ were aligned on the structure of the C nonamer that was already available in the PDB (PDB: 7ALW^36^). The obtained models for the full protein complex were relaxed by OpenMM, resulting in a model with very high global accuracy (TM-score=0.98), but slightly less accurate interfaces (ICS=0.45) (Fig. 6B).

H1114 was a large heteromeric complex with stoichiometry A_4_B_8_C_8_. Using the PPI3D server we identified templates for the B_2_C_2_ subcomplex. AlphaFold produced a confident model for this subcomplex, which was in complete agreement with these structural templates. The A_4_ tetramers were also modeled consistently using AlphaFold. In the next step, we modeled several variants of subcomplexes composed of either A and B subunits or all three subunits. One of the resulting subcomplexes, A_4_B_2_, was further used as a core to align all the subcomplexes and to get the final assembly model of high accuracy (ICS > 0.8 and TM-score > 0.85) (Fig. 6C). A similar combination of partial AlphaFold models allowed us to obtain a fairly accurate model for H1137, a large heteromeric protein complex of 10 chains (ABCDEFG_2_HI) related to lipid transport across the bacterial cell membrane (ICS=0.82, TM-score=0.80) (Fig. 6D). In this case the entire protein complex could be modeled by AlphaFold, but the model assembled from partial AlphaFold models was more accurate.

### Modeling of antibody-antigen interactions using AlphaFold and docking

The antibody-antigen complexes are a special case of protein interactions. Different antibodies can bind to different sites of the surface of the same antigen^37^, thus neither the homology-based nor the coevolution-based modeling methods are guaranteed to produce reliable models. Therefore, it is not surprising that AlphaFold-Multimer is often unable to correctly model the antibody-antigen complexes^6, 7^. CASP15 had 8 such targets (5 nanobody-CNPase complexes and 3 antibody-SARS-CoV-2 capsid protein complexes). Being aware that for AlphaFold such interactions are challenging, we additionally used rigid-body docking for these targets to obtain large and diverse sets of models. Interestingly, AlphaFold and docking methods either both worked or both failed on specific antibody-antigen targets (Supplementary Figure S6). At first glance, our failures in modeling antibody-antigen interactions (H1142, H1144, H1166v1, H1167v1) can be attributed to insufficient sampling (Supplementary Table S2).

Several previous studies have shown that the accuracy of protein-protein docking models depends largely on the monomer quality, so that even small inconsistencies may result in incorrect predictions^28, 33, 38^. The antibodies recognize the antigens using the hypervariable complementarity-determining region (CDR) loops. The main issue in antibody structure prediction is the modeling of the CDRs, especially the CDR-H3 loop^39, 40^, and the conformations of these loops may even change upon binding^41^. Indeed, our failures to predict the antibody-antigen complexes seem to be related to incorrect positioning of CDR loops in the antibody structures (Supplementary Figure S7). Thus, it appears that a more aggressive sampling at the stage of modeling an antibody-antigen complex or an increased sampling of antibody monomers prior to docking might be a way to improve modeling success for this type of complexes.

## Discussion

CASP15 revealed that even a standard AlphaFold-Multimer pipeline is able to produce accurate structures for a significant fraction of protein complexes. Analysis of our results showed that by forcing AlphaFold-Multimer to do a more extensive conformational sampling, the standard performance can be increased even further. In our case, a more extensive sampling was achieved by providing custom MSAs with different pairing strategies. Enabling dropout in the neural network^42^ or using other ways to increase sampling without retraining the main neural network model^6^ might be some of the alternative strategies. What has also become clear from our CASP15 results is that template-based approaches for modeling protein assemblies are no match to AlphaFold. Thus, template-based models may serve as a ‘quick-and-dirty’ way to get an approximate model, but if the details of a protein complex are important, homology modeling is no longer the method of choice.

Although the use of AlphaFold led to an impressive advancement in modeling protein assemblies, challenges remain. Consistent with previous observations^6–8^, our results showed that two types of protein complexes, namely, large multisubunit assemblies and antibody-antigen complexes, are particularly challenging for AlphaFold. As we showed here, large complexes could be successfully modeled by hybrid approaches such as combining overlapping AlphaFold-generated subcomplexes or docking of monomers/non-overlapping subcomplexes followed by the interface-focused scoring. Analysis of our models for antibody-antigen complexes suggests that the likely reason for failures is insufficient sampling of antibody CDR conformations. Although aggressive sampling by AlphaFold coupled with self-assessment^42^ might be successful in some cases, it is unlikely to completely solve the problem. Perhaps a more universal solution might be an extensive sampling of antigen CDR conformations followed by antibody-antigen docking. CDR modeling/sampling may well be performed by other promising methods such as OmegaFold, in a preliminary report presented as an accurate predictor of antibody loops^43^.

A scoring method, independent of intrinsic accuracy estimates of a given structure prediction method (e.g., AlphaFold), is certainly needed when hybrid techniques are used as exemplified by our protocol. Such a scoring method is also essential if structural models are generated by different approaches, because no matter how good an individual modeling approach is in self-assessment, an independent “judge” is needed to decide on the best model. Already a number of other methods such as RoseTTAFold^44^, OmegaFold^43^ and ESMFold^45^ may be used to complement AlphaFold. Therefore, the importance of independent scoring methods might be expected to increase as the number of diverse methods for structure prediction continues to grow.

## Software availability

https://github.com/kliment-olechnovic/ftdmp

## Acknowledgments

We wish to thank the CASP15 organizers and CASP and CAPRI assessors as well as the experimentalists who provided unpublished 3D structures of protein assemblies as prediction targets.

## References

1. Kryshtafovych A, Schwede T, Topf M, Fidelis K, Moult J. Critical assessment of methods of protein structure prediction (CASP)-Round XIV. Proteins 2021;89(12):1607–1617.

2. Jumper J, Evans R, Pritzel A, Green T, Figurnov M, Ronneberger O, Tunyasuvunakool K, Bates R, Žídek A, Potapenko A, Bridgland A, Meyer C, Kohl SAA, Ballard AJ, Cowie A, Romera-Paredes B, Nikolov S, Jain R, Adler J, Back T, Petersen S, Reiman D, Clancy E, Zielinski M, Steinegger M, Pacholska M, Berghammer T, Bodenstein S, Silver D, Vinyals O, Senior AW, Kavukcuoglu K, Kohli P, Hassabis D. Highly accurate protein structure prediction with AlphaFold. Nature 2021;596(7873):583–589.

3. Tunyasuvunakool K, Adler J, Wu Z, Green T, Zielinski M, Žídek A, Bridgland A, Cowie A, Meyer C, Laydon A, Velankar S, Kleywegt GJ, Bateman A, Evans R, Pritzel A, Figurnov M, Ronneberger O, Bates R, Kohl SAA, Potapenko A, Ballard AJ, Romera-Paredes B, Nikolov S, Jain R, Clancy E, Reiman D, Petersen S, Senior AW, Kavukcuoglu K, Birney E, Kohli P, Jumper J, Hassabis D. Highly accurate protein structure prediction for the human proteome. Nature 2021;596(7873):590– 596.

4. Varadi M, Anyango S, Deshpande M, Nair S, Natassia C, Yordanova G, Yuan D, Stroe O, Wood G, Laydon A, Žídek A, Green T, Tunyasuvunakool K, Petersen S, Jumper J, Clancy E, Green R, Vora A, Lutfi M, Figurnov M, Cowie A, Hobbs N, Kohli P, Kleywegt G, Birney E, Hassabis D, Velankar S. AlphaFold Protein Structure Database: massively expanding the structural coverage of protein-sequence space with high-accuracy models. Nucleic Acids Res 2022;50(D1):D439–D444.

5. Mirdita M, Schütze K, Moriwaki Y, Heo L, Ovchinnikov S, Steinegger M. ColabFold: making protein folding accessible to all. Nat Methods 2022;19(6):679–682.

6. Evans R, O’Neill M, Pritzel A, Antropova N, Senior A, Green T, Žídek dek A, Bates R, Blackwell S, Yim J, Ronneberger O, Bodenstein S, Zielinski M, Bridgland A, Potapenko A, Cowie A, Tunyasuvunakool K, Jain R, Clancy E, Kohli P, Jumper J, Hassabis D. Protein complex prediction with AlphaFold-Multimer. bioRxiv 2022:2021.10.04.463034.

7. Yin R, Feng BY, Varshney A, Pierce BG. Benchmarking AlphaFold for protein complex modeling reveals accuracy determinants. Protein Sci 2022;31(8):e4379.

8. Bryant P, Pozzati G, Zhu W, Shenoy A, Kundrotas P, Elofsson A. Predicting the structure of large protein complexes using AlphaFold and Monte Carlo tree search. Nat Commun 2022;13(1):6028.

9. Eddy SR. Accelerated Profile HMM Searches. PLoS Comput Biol 2011;7(10):e1002195.

10. Roux S, Páez-Espino D, Chen I-MA, Palaniappan K, Ratner A, Chu K, Reddy TBK, Nayfach S, Schulz F, Call L, Neches RY, Woyke T, Ivanova NN, Eloe-Fadrosh EA, Kyrpides NC. IMG/VR v3: an integrated ecological and evolutionary framework for interrogating genomes of uncultivated viruses. Nucleic Acids Res 2021;49(D1):D764–D775.

11. Mitchell AL, Almeida A, Beracochea M, Boland M, Burgin J, Cochrane G, Crusoe MR, Kale V, Potter SC, Richardson LJ, Sakharova E, Scheremetjew M, Korobeynikov A, Shlemov A, Kunyavskaya O, Lapidus A, Finn RD. MGnify: the microbiome analysis resource in 2020. Nucleic Acids Res 2020;48(D1):D570–D578.

12. UniProt Consortium. UniProt: the Universal Protein Knowledgebase in 2023. Nucleic Acids Res 2023;51(D1):D523–D531.

13. O’Leary NA, Wright MW, Brister JR, Ciufo S, Haddad D, McVeigh R, Rajput B, Robbertse B, Smith-White B, Ako-Adjei D, Astashyn A, Badretdin A, Bao Y, Blinkova O, Brover V, Chetvernin V, Choi J, Cox E, Ermolaeva O, Farrell CM, Goldfarb T, Gupta T, Haft D, Hatcher E, Hlavina W, Joardar VS, Kodali VK, Li W, Maglott D, Masterson P, McGarvey KM, Murphy MR, O’Neill K, Pujar S, Rangwala SH, Rausch D, Riddick LD, Schoch C, Shkeda A, Storz SS, Sun H, Thibaud-Nissen F, Tolstoy I, Tully RE, Vatsan AR, Wallin C, Webb D, Wu W, Landrum MJ, Kimchi A, Tatusova T, DiCuccio M, Kitts P, Murphy TD, Pruitt KD. Reference sequence (RefSeq) database at NCBI: current status, taxonomic expansion, and functional annotation. Nucleic Acids Res 2016;44(D1):D733–745.

14. Gabb HA, Jackson RM, Sternberg MJ. Modelling protein docking using shape complementarity, electrostatics and biochemical information. J Mol Biol 1997;272(1):106–120.

15. Ritchie DW, Kemp GJ. Protein docking using spherical polar Fourier correlations. Proteins 2000;39(2):178–194.

16. Ritchie DW, Grudinin S. Spherical polar Fourier assembly of protein complexes with arbitrary point group symmetry. J Appl Cryst 2016;49(1):158–167.

17. Kryshtafovych A, Moult J, Billings WM, Della Corte D, Fidelis K, Kwon S, Olechnovič K, Seok C, Venclovas Č, Won J, CASP-COVID participants. Modeling SARS-CoV-2 proteins in the CASP-commons experiment. Proteins 2021;89(12):1987–1996.

18. Olechnovič K, Venclovas Č. VoroMQA: Assessment of protein structure quality using interatomic contact areas. Proteins 2017;85(6):1131–1145.

19. Olechnovič K, Venclovas Č. Contact Area-Based Structural Analysis of Proteins and Their Complexes Using CAD-Score. Methods Mol Biol 2020;2112:75–90.

20. Eastman P, Swails J, Chodera JD, McGibbon RT, Zhao Y, Beauchamp KA, Wang L-P, Simmonett AC, Harrigan MP, Stern CD, Wiewiora RP, Brooks BR, Pande VS. OpenMM 7: Rapid development of high performance algorithms for molecular dynamics. PLoS Comput Biol 2017;13(7):e1005659.

21. Zimmermann L, Stephens A, Nam S-Z, Rau D, Kübler J, Lozajic M, Gabler F, Söding J, Lupas AN, Alva V. A Completely Reimplemented MPI Bioinformatics Toolkit with a New HHpred Server at its Core. J Mol Biol 2018;430(15):2237–2243.

22. Margelevičius M. COMER2: GPU-accelerated sensitive and specific homology searches. Bioinformatics 2020;36(11):3570–3572.

23. Dapkūnas J, Margelevičius M. The COMER web server for protein analysis by homology. Bioinformatics 2023;39(1):btac807.

24. Holm L. Dali server: structural unification of protein families. Nucleic Acids Res 2022;50(W1):W210–W215.

25. Dapkūnas J, Timinskas A, Olechnovič K, Margelevičius M, Dičiūnas R, Venclovas Č. The PPI3D web server for searching, analyzing and modeling protein-protein interactions in the context of 3D structures. Bioinformatics 2017;33(6):935–937.

26. Burley SK, Bhikadiya C, Bi C, Bittrich S, Chen L, Crichlow GV, Christie CH, Dalenberg K, Di Costanzo L, Duarte JM, Dutta S, Feng Z, Ganesan S, Goodsell DS, Ghosh S, Green RK, Guranović V, Guzenko D, Hudson BP, Lawson CL, Liang Y, Lowe R, Namkoong H, Peisach E, Persikova I, Randle C, Rose A, Rose Y, Sali A, Segura J, Sekharan M, Shao C, Tao Y-P, Voigt M, Westbrook JD, Young JY, Zardecki C, Zhuravleva M. RCSB Protein Data Bank: powerful new tools for exploring 3D structures of biological macromolecules for basic and applied research and education in fundamental biology, biomedicine, biotechnology, bioengineering and energy sciences. Nucleic Acids Res 2021;49(D1):D437–D451.

27. Dapkūnas J, Venclovas Č. Template-Based Modeling of Protein Complexes Using the PPI3D Web Server. Methods Mol Biol 2020;2165:139–155.

28. Dapkūnas J, Olechnovič K, Venclovas Č. Modeling of protein complexes in CASP14 with emphasis on the interaction interface prediction. Proteins 2021;89(12):1834–1843.

29. Jones DT, Cozzetto D. DISOPRED3: precise disordered region predictions with annotated protein-binding activity. Bioinformatics 2015;31(6):857–863.

30. Lafita A, Bliven S, Kryshtafovych A, Bertoni M, Monastyrskyy B, Duarte JM, Schwede T, Capitani G. Assessment of protein assembly prediction in CASP12. Proteins 2018;86 Suppl 1:247–256.

31. Mariani V, Biasini M, Barbato A, Schwede T. lDDT: a local superposition-free score for comparing protein structures and models using distance difference tests. Bioinformatics 2013;29(21):2722– 2728.

32. Zhang C, Shine M, Pyle AM, Zhang Y. US-align: universal structure alignments of proteins, nucleic acids, and macromolecular complexes. Nat Methods 2022;19(9):1109–1115.

33. Dapkūnas J, Olechnovič K, Venclovas Č. Structural modeling of protein complexes: Current capabilities and challenges. Proteins 2019;87(12):1222–1232.

34. Ozden B, Kryshtafovych A, Karaca E. Assessment of the CASP14 assembly predictions. Proteins 2021;89(12):1787–1799.

35. Lensink MF, Brysbaert G, Mauri T, Nadzirin N, Velankar S, Chaleil RAG, Clarence T, Bates PA, Kong R, Liu B, Yang G, Liu M, Shi H, Lu X, Chang S, Roy RS, Quadir F, Liu J, Cheng J, Antoniak A, Czaplewski C, Giełdoń a, Kogut M, Lipska AG, Liwo A, Lubecka EA, Maszota-Zieleniak M, Sieradzan AK, Ślusarz R, Wesołowski PA, Zięba K, Del Carpio Muñoz CA, Ichiishi E, Harmalkar A, Gray JJ, Bonvin Amjj, Ambrosetti F, Vargas Honorato R, Jandova Z, Jiménez-García B, Koukos PI, Van Keulen S, Van Noort CW, Réau M, Roel-Touris J, Kotelnikov S, Padhorny D, Porter KA, Alekseenko A, Ignatov M, Desta I, Ashizawa R, Sun Z, Ghani U, Hashemi N, Vajda S, Kozakov D, Rosell M, Rodríguez-Lumbreras LA, Fernandez-Recio J, Karczynska A, Grudinin S, Yan Y, Li H, Lin P, Huang S-Y, Christoffer C, Terashi G, Verburgt J, Sarkar D, Aderinwale T, Wang X, Kihara D, Nakamura T, Hanazono Y, Gowthaman R, Guest JD, Yin R, Taherzadeh G, Pierce BG, Barradas-Bautista D, Cao Z, Cavallo L, Oliva R, Sun Y, Zhu S, Shen Y, Park T, Woo H, Yang J, Kwon S, Won J, Seok C, Kiyota Y, Kobayashi S, Harada Y, Takeda-Shitaka M, Kundrotas PJ, Singh A, Vakser IA, Dapkūnas J, Olechnovič K, Venclovas Č, Duan R, Qiu L, Xu X, Zhang S, Zou X, Wodak SJ. Prediction of protein assemblies, the next frontier: The CASP14-CAPRI experiment. Proteins 2021;89(12):1800–1823.

36. Kuhlen L, Johnson S, Cao J, Deme JC, Lea SM. Nonameric structures of the cytoplasmic domain of FlhA and SctV in the context of the full-length protein. PLoS One 2021;16(6):e0252800.

37. Sela-Culang I, Kunik V, Ofran Y. The structural basis of antibody-antigen recognition. Front Immunol 2013;4:302.

38. Singh A, Dauzhenka T, Kundrotas PJ, Sternberg MJE, Vakser IA. Application of docking methodologies to modeled proteins. Proteins 2020;88(9):1180–1188.

39. Norman RA, Ambrosetti F, Bonvin Amjj, Colwell LJ, Kelm S, Kumar S, Krawczyk K. Computational approaches to therapeutic antibody design: established methods and emerging trends. Brief Bioinform 2020;21(5):1549–1567.

40. Hummer AM, Abanades B, Deane CM. Advances in computational structure-based antibody design. Curr Opin Struct Biol 2022;74:102379.

41. Guest JD, Vreven T, Zhou J, Moal I, Jeliazkov JR, Gray JJ, Weng Z, Pierce BG. An expanded benchmark for antibody-antigen docking and affinity prediction reveals insights into antibody recognition determinants. Structure 2021;29(6):606-621.e5.

42. Wallner B. AFsample: Improving Multimer Prediction with AlphaFold using Aggressive Sampling. bioRxiv; 2022. p 2022.12.20.521205.

43. Wu R, Ding F, Wang R, Shen R, Zhang X, Luo S, Su C, Wu Z, Xie Q, Berger B, Ma J, Peng J. High-resolution de novo structure prediction from primary sequence. bioRxiv; 2022. p 2022.07.21.500999.

44. Baek M, DiMaio F, Anishchenko I, Dauparas J, Ovchinnikov S, Lee GR, Wang J, Cong Q, Kinch LN, Schaeffer RD, Millán C, Park H, Adams C, Glassman CR, DeGiovanni A, Pereira JH, Rodrigues AV, Dijk AA van, Ebrecht AC, Opperman DJ, Sagmeister T, Buhlheller C, Pavkov-Keller T, Rathinaswamy MK, Dalwadi U, Yip CK, Burke JE, Garcia KC, Grishin NV, Adams PD, Read RJ, Baker D. Accurate prediction of protein structures and interactions using a three-track neural network. Science 2021;373(6557):871–876.

45. Lin Z, Akin H, Rao R, Hie B, Zhu Z, Lu W, Smetanin N, Verkuil R, Kabeli O, Shmueli Y, Costa A dos S, Fazel-Zarandi M, Sercu T, Candido S, Rives A. Evolutionary-scale prediction of atomic level protein structure with a language model. bioRxiv; 2022. p 2022.07.20.500902.

