## Supplementary data for "Prediction of protein assemblies by structure sampling followed by interface-focused scoring"

#### Supplementary information

**Corresponding author**:

Česlovas Venclovas
Institute of Biotechnology, Life Sciences Center
Vilnius University
Saulėtekio 7,
LT-10257 Vilnius, Lithuania

### Supplementary Methods

#### VoroIF-jury algorithm

Input**:**

- multimeric structural models of the same target
- target stoichiometry
- target chain sequences
- N (maximum number of top models to consider from each QA ranking)

Output**:**

- model names ordered by the model quality estimates (VoroIF-jury scores)

Algorithm steps:

1. Collect all the available structural models (possibly from diverse sources, e.g. AlphaFold-Multimer, docking, template-based modeling).
2. Using the target stoichiometry and chain sequences, automatically fix chain naming and sequence numbering in all the models.
3. Cluster all the models using the interface CAD-score, select cluster representatives to analyze further.
4. Compute multiple QA rankings using different interface-focused scores (variations of VoroMQA-energy, VoroMQA-light, VoroMQA-dark, and VoroIF-GNN scores). The list of rankings is presented after the algorithm.
5. For every X from 1 to N:
    a. Select top X models from every available QA ranking.
    b. Pool all the selected models into a superset (if a model was selected by more than one QA method, it is included multiple times, thus popular models gain more weight).
    c. For every model in the superset, calculate the average of all its pairwise similarities (interface CAD-score values) with the other models in the superset. This value is called "top X interface consensus value".
6. For every model, select the highest achieved top X interface consensus value (among all X from 1 to N). This value is called "VoroIF-jury score".
7. Return the list of model names ordered by their VoroIF-jury score (from highest to lowest).
8. (optionally) Diversify the final ranking so that every model is not too similar to any of the higher ranked ones.

**QA rankings used in the VoroIF-jury algorithm**

- ranking based on the total **VoroMQA-energy** of the inter-chain interface
- tournament-based ranking using the total interface **VoroMQA-energy** and the interface **clash score**
- ranking based on the the total **generic VoroMQA-energy** of the inter-chain interface
- tournament-based ranking using the total interface **generic VoroMQA-energy** and the interface **clash score**
- ranking based on the sum of **VoroIF-GNN** scores of the inter-chain interface contacts
- ranking based on the normalized interface **VoroIF-GNN** score (the sum of **VoroIF-GNN** scores of the interface contacts divided by the total interface area)
- tournament-based ranking using the sum of interface **VoroIF-GNN** scores and the normalized interface **VoroIF-GNN** score
- ranking based on the average normalized **VoroIF-GNN** score per interface residue
- ranking based on the weighted average normalized **VoroIF-GNN** score per interface residue (residue weight is the total area of that residue inter-chain contacts)
- tournament-based ranking using the sum of interface **VoroIF-GNN** scores and the interface **clash score**
- tournament-based ranking using the sum of interface **VoroIF-GNN** scores and the total interface **VoroMQA-energy**
- tournament-based ranking using the normalized interface **VoroIF-GNN** score and the total interface **VoroMQA-energy**
- tournament-based ranking using the the weighted average normalized per-residue **VoroIF-GNN** score and the total interface **VoroMQA-energy**
- tournament-based ranking using the normalized interface **VoroIF-GNN** score and the total generic interface **VoroMQA-energy**
- tournament-based ranking using the the average normalized per-residue **VoroIF-GNN** score and the total interface **generic VoroMQA-energy**
- tournament-based ranking using the the weighted average normalized per-residue **VoroIF-GNN** score and the total interface **generic VoroMQA-energy**
- tournament-based ranking using the the total interface **VoroMQA-energy** and the **VoroMQA-light** global score (the same as VoroMQA-select-2018)
- tournament-based ranking using the the total interface **VoroMQA-energy** and the **VoroMQA-dark** global score (the same as VoroMQA-select-2020)
- tournament-based ranking using the the sum of interface **VoroIF-GNN** scores and the **VoroMQA-light** global score
- tournament-based ranking using the the sum of interface **VoroIF-GNN** scores and the **VoroMQA-dark** global score

#### Supplementary Tables

**Supplementary Table S1.** The accuracy of our most confident models (model 1) for the CASP15 targets

| Target | Stoichiometry | Difficulty | ICS | IPS | lDDT oligo | TM-score |
|---|---|---|---|---|---|---|
| H1106 | AB | Easy | 0.67 | 0.67 | 0.80 | 0.85 |
| T1109 | $A_2$ | Easy | 0.82 | 0.86 | 0.87 | 0.96 |
| T1110 | $A_2$ | Easy | 0.91 | 0.95 | 0.92 | 0.98 |
| H1111 | $A_9B_9C_9$ | Medium | 0.45 | 0.42 | 0.73 | 0.98 |
| T1113 | $A_2$ | Hard | 0.90 | 0.90 | 0.91 | 0.93 |
| H1114 | $A_4B_8B_8$ | Medium | 0.81 | 0.82 | 0.87 | 0.94 |
| H1114v2 | $A_4B_2$ | Medium | 0.61 | 0.55 | 0.84 | 0.73 |
| T1115 | $A_{16}$ | Medium | 0.68 | 0.70 | 0.69 | 0.90 |
| T1121 | $A_2$ | Medium | 0.41 | 0.25 | 0.77 | 0.55 |
| T1123 | $A_2$ | Medium | 0.30 | 0.56 | 0.56 | 0.81 |
| T1124 | $A_2$ | Easy | 0.83 | 0.81 | 0.86 | 0.96 |
| T1127 | $A_2$ | Easy | 0.88 | 0.91 | 0.88 | 0.98 |
| H1129 | AB | Medium | 0.75 | 0.83 | 0.82 | 0.97 |
| T1132 | $A_6$ | Medium | 0.86 | 0.89 | 0.92 | 0.99 |
| H1134 | AB | Medium | 0.76 | 0.79 | 0.89 | 0.97 |
| H1135 | $A_9B_3$ | Medium | 0.60 | 0.63 | 0.78 | 0.51 |
| H1137 | $ABCDEFG_2HI$ | Medium | 0.82 | 0.84 | 0.80 | 0.80 |
| H1140 | AB | Medium | 0.27 | 0.47 | 0.75 | 0.73 |
| H1141 | AB | Medium | 0.72 | 0.69 | 0.84 | 0.96 |
| H1142 | AB | Medium | 0.00 | 0.13 | 0.79 | 0.63 |
| H1143 | AB | Medium | 0.83 | 0.80 | 0.87 | 0.98 |
| H1144 | AB | Medium | 0.00 | 0.11 | 0.78 | 0.62 |
| H1151 | AB | Easy | 0.73 | 0.69 | 0.84 | 0.95 |
| T1153 | $A_2$ | Medium | 0.81 | 0.76 | 0.87 | 0.93 |
| H1157 | AB | Medium | 0.72 | 0.66 | 0.70 | 0.79 |
| T1160 | $A_2$ | Easy | 0.15 | 0.48 | 0.44 | 0.34 |
| T1161 | $A_2$ | Easy | 0.41 | 0.72 | 0.53 | 0.59 |
| H1166v1 | ABC | Medium | 0.00 | 0.08 | 0.79 | 0.64 |
| H1167v1 | ABC | Medium | 0.00 | 0.24 | 0.83 | 0.51 |
| H1168v1 | ABC | Medium | 0.79 | 0.79 | 0.89 | 0.86 |
| T1170 | $A_6$ | Easy | 0.57 | 0.63 | 0.81 | 0.93 |
| H1171 | $A_6B$ | Medium | 0.58 | 0.61 | 0.78 | 0.91 |
| H1172 | $A_6B_2$ | Medium | 0.60 | 0.64 | 0.77 | 0.90 |
| T1173 | $A_3$ | Medium | 0.55 | 0.50 | 0.61 | 0.63 |
| T1174 | $A_3$ | Hard | 0.86 | 0.86 | 0.86 | 0.73 |
| T1176 | $A_8$ | Medium | 0.01 | 0.08 | 0.47 | 0.24 |
| T1176v1 | $A_2$ | Medium | 0.00 | 0.05 | 0.52 | 0.41 |
| T1178 | $A_2$ | Easy | 0.60 | 0.76 | 0.78 | 0.93 |
| T1179 | $A_2$ | Easy | 0.12 | 0.44 | 0.61 | 0.61 |
| T1181 | $A_3$ | Easy | 0.46 | 0.52 | 0.74 | 0.43 |
| H1185 | ABCD | Easy | 0.69 | 0.63 | 0.79 | 0.97 |
| T1187 | $A_2$ | Hard | 0.00 | 0.17 | 0.77 | 0.49 |
| T1192 | $A_{10}$ | Easy | 0.71 | 0.62 | 0.85 | 0.88 |

.

**Supplementary Table S2.** Maximum available interface CAD-score values for the models of failed targets

| Target | Our 'model 1' | Best of our five submitted models | Best of our all generated models | Best of all CASP15 models |
|---|---|---|---|---|
| H1142 | 0.00 | 0.00 | 0.10 | 0.27 |
| H1144 | 0.00 | 0.06 | 0.18 | 0.66 |
| T1160 | 0.14 | 0.23 | 0.24 | 0.63 |
| H1166v1 | 0.00 | 0.02 | 0.12 | 0.21 |
| H1167v1 | 0.00 | 0.04 | 0.10 | 0.12 |
| T1176 | 0.01 | 0.01 | 0.01 | 0.03 |
| T1176v1 | 0.01 | 0.01 | 0.01 | 0.01 |
| T1179 | 0.09 | 0.10 | 0.16 | 0.71 |
| T1187 | 0.00 | 0.74 | 0.74 | 0.75 |

**Supplementary Table S3.** Comparison of accuracy scores of our template-based and AlphaFold models

| Target | Best template-based model | | | Best AlphaFold model | | |
|---|---|---|---|---|---|---|
| | Name | ICS | TM-score | Name | ICS | TM-score |
| T1109 | T1109TS494_5o | 0.44 | 0.85 | T1109TS494_1o | 0.82 | 0.96 |
| T1110 | T1110TS494_4o | 0.79 | 0.88 | T1110TS494_1o | 0.91 | 0.98 |
| T1121 | T1121TS494_3o | 0.07 | 0.30 | T1121TS494_1o | 0.41 | 0.55 |
| T1121 | T1121TS494_3o | 0.07 | 0.30 | T1121TS494_4o | 0.55 | 0.72 |
| T1123 | T1123TS494_5o | 0.04 | 0.35 | T1123TS494_1o | 0.30 | 0.81 |
| T1123 | T1123TS494_5o | 0.04 | 0.35 | T1123TS494_3o | 0.43 | 0.91 |
| T1124 | T1124TS494_5o | 0.25 | 0.55 | T1124TS494_1o | 0.83 | 0.96 |
| T1127 | T1127TS494_5o | 0.65 | 0.70 | T1127TS494_1o | 0.88 | 0.98 |
| T1132 | T1132TS494_4o | 0.47 | 0.50 | T1132TS494_1o | 0.86 | 0.99 |
| H1137 | H1137TS494_5 | 0.19 | 0.40 | H1137TS494_3 | 0.70 | 0.68 |
| H1151 | H1151TS494_5 | 0.61 | 0.74 | H1151TS494_1 | 0.73 | 0.95 |
| T1153 | T1153TS494_3o | 0.60 | 0.40 | T1153TS494_1o | 0.81 | 0.93 |
| T1160 | T1160TS494_2o | 0.15 | 0.27 | T1160TS494_1o | 0.15 | 0.34 |
| T1161 | T1161TS494_3o | 0.32 | 0.46 | T1161TS494_1o | 0.41 | 0.59 |
| H1166v1 | H1166v1TS494_3 | 0.00 | 0.65 | H1166v1TS494_4 | 0.00 | 0.75 |
| H1167v1 | H1167v1TS494_4 | 0.00 | 0.60 | H1167v1TS494_5 | 0.10 | 0.54 |
| H1168v1 | H1168v1TS494_4 | 0.61 | 0.96 | H1168v1TS494_1 | 0.79 | 0.86 |
| T1170 | T1170TS494_4o | 0.45 | 0.86 | T1170TS494_1o | 0.57 | 0.93 |
| T1179 | T1179TS494_3o | 0.19 | 0.64 | T1179TS494_4o | 0.10 | 0.77 |
| T1179 | T1179TS494_3o | 0.19 | 0.64 | T1179TS494_5o | 0.22 | 0.90 |
| T1181 | T1181TS494_5o | 0.04 | 0.36 | T1181TS494_1o | 0.46 | 0.43 |
| T1181 | T1181TS494_5o | 0.04 | 0.36 | T1181TS494_2o | 0.53 | 0.84 |
| H1185 | H1185TS494_4 | 0.24 | 0.70 | H1185TS494_1 | 0.69 | 0.97 |

#### Supplementary Figures

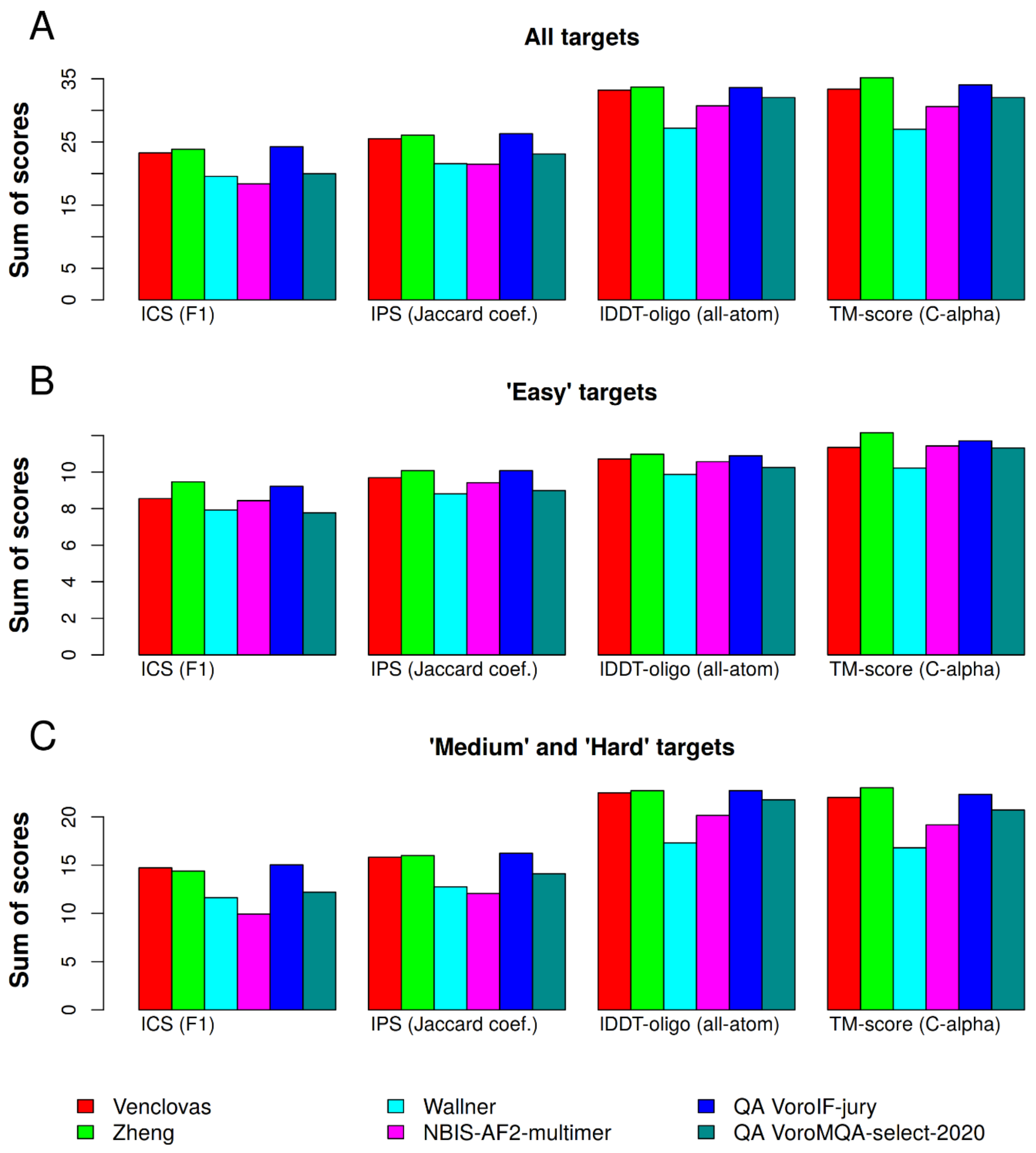


**Supplementary Figure S1**. Comparison of results of our group ("Venclovas") and our automated model selection protocols (the older "VoroMQA-select-2020" and the latest "VoroIF-jury) with other two top-performing CASP15 groups ("Zheng" and "Wallner") and with the baseline group that run the standard AlphaFold-Multimer ("NBIS-AF2-multimer"). The sums of scores are presented for the three subsets of targets defined using the target dufficulty classification done by the CASP15 assessors: all the 43 targets (A), 14 "Easy" targets (B), 29 "Medium" or "Hard" targets (C)

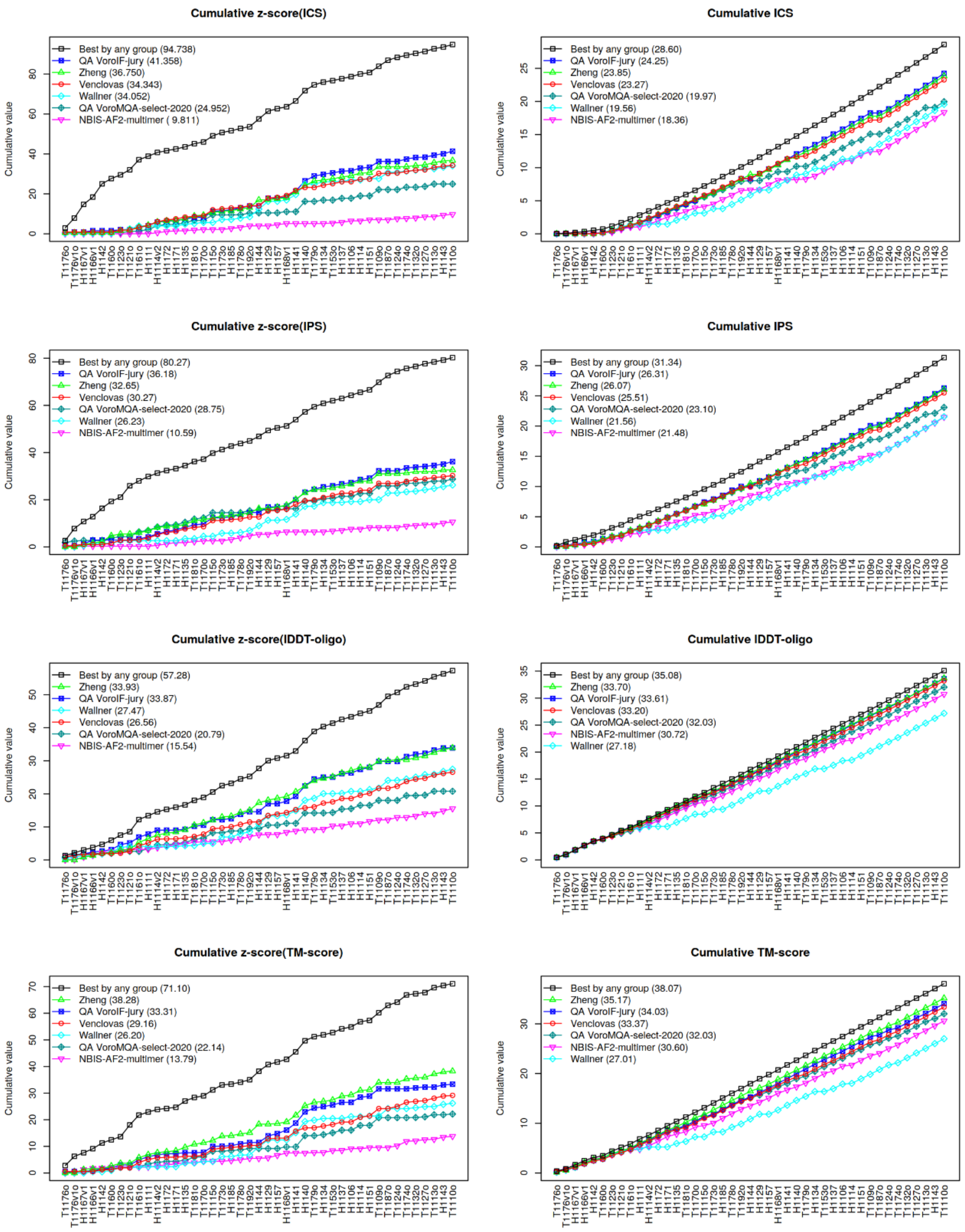


**Supplementary Figure S2**. Cumulative z-score (left side) and raw score (right side) values of the models designated as first. Targets were ordered by the maximum achieved ICS score. The group names in the plot legend are sorted by the corresponding achieved sum (the sum values are shown in brackets)

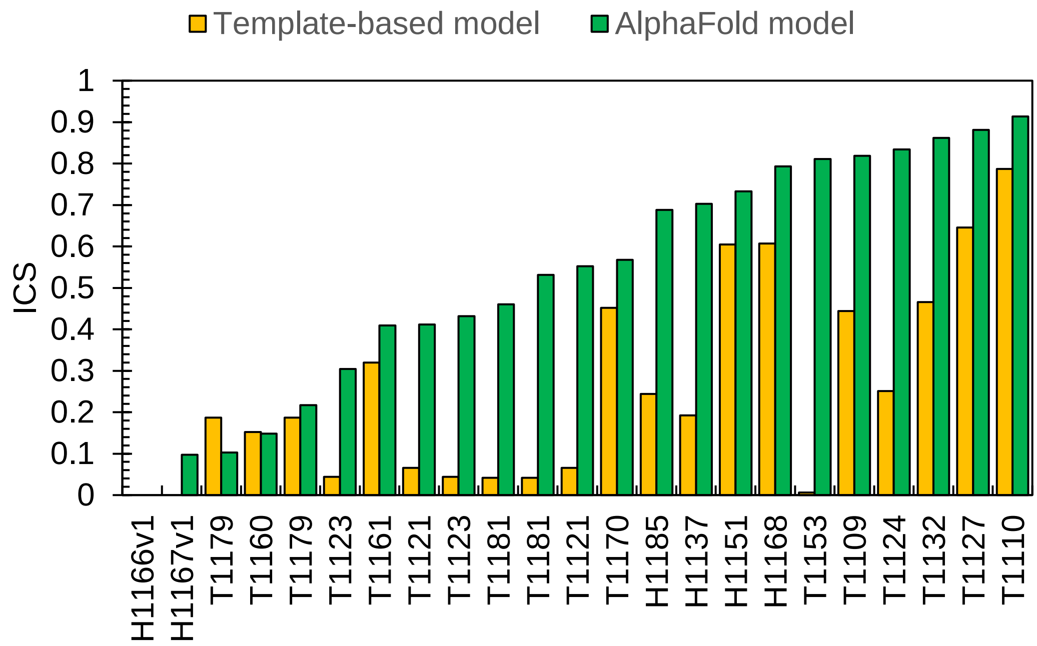


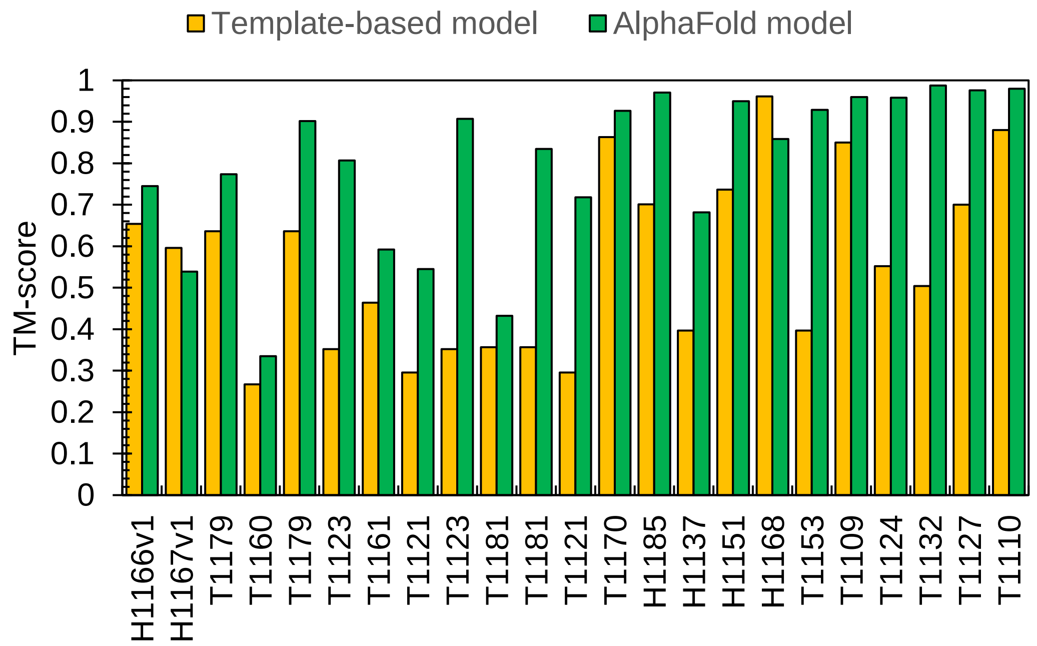


**Supplementary Figure S3**. Comparison of our template-based and AlphaFold model accuracy for CASP15 targets, for which structural templates were identified. Targets are sorted by the highest ICS of AlphaFold models

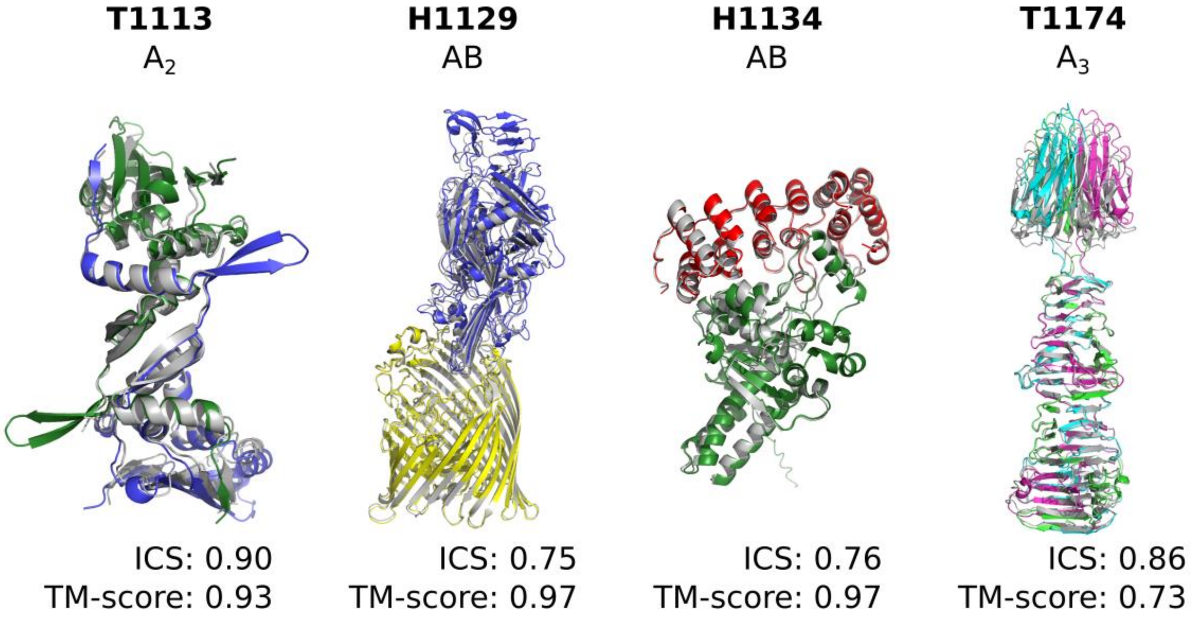


**Supplementary Figure S4**. Examples of accurate models produced by AlphaFold-Multimer despite the absence of multimeric templates in the PDB. Experimental structures are colored gray, model chains are shown in different colors. Models were generated using the ColabFold pipeline and custom MSAs

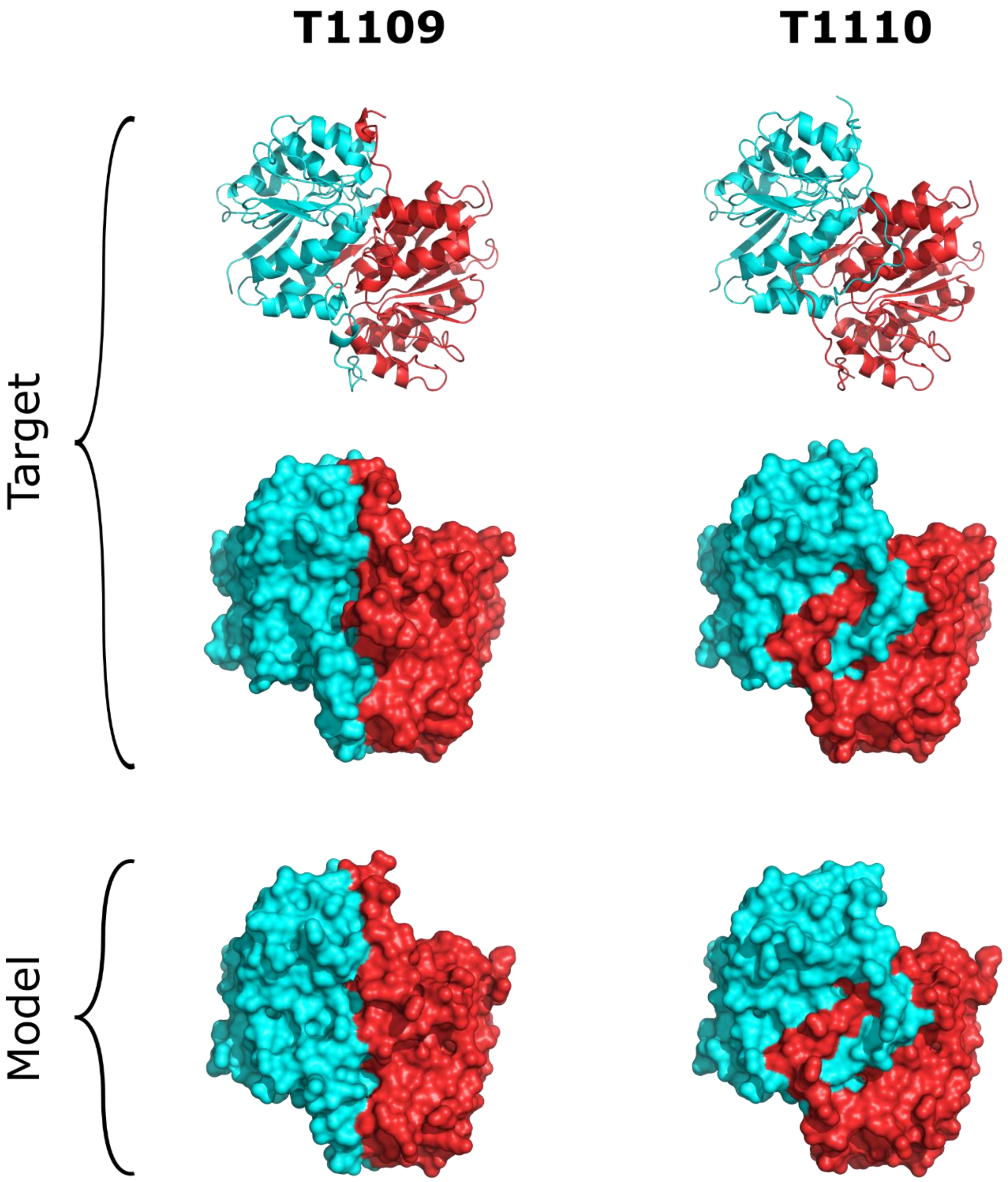


**Supplementary Figure S5**. Alternative conformations of mutated (T1109) and wild-type (T1110) isocyanide hydratase from *Ralstonia solanacearum*

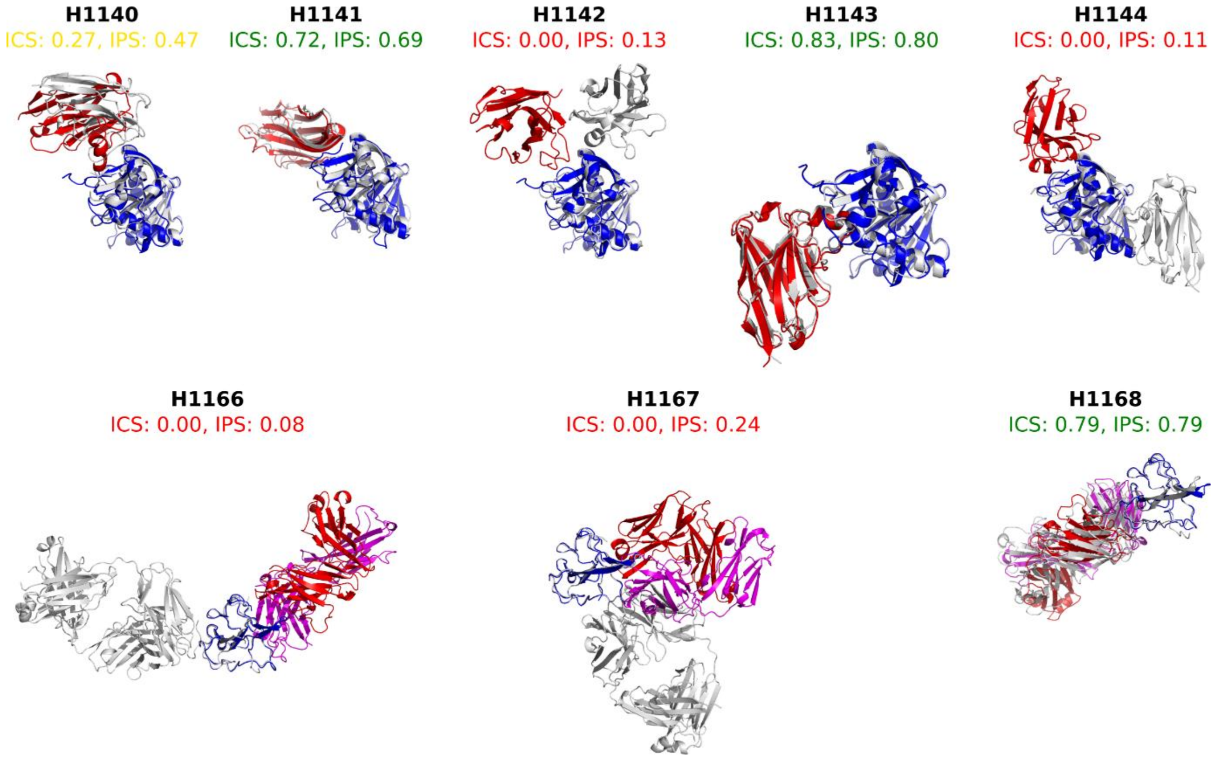


**Supplementary Figure S6**. Modeling results for CASP15 antibody-antigen interaction targets. Experimental structures are shown in gray, antigens are blue, and antibodies are colored in red and magenta

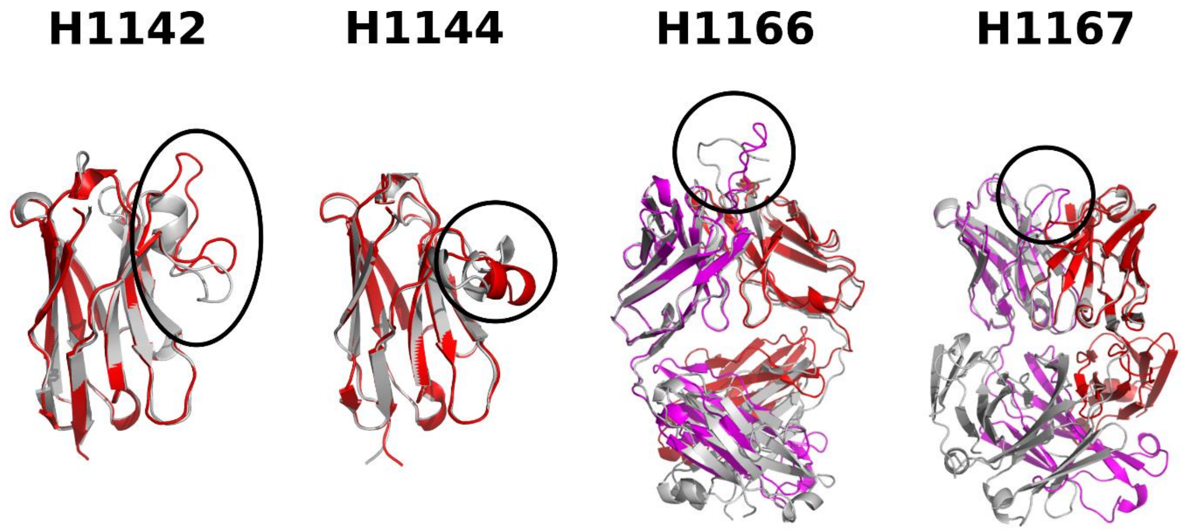


**Supplementary Figure S7**. Incorrect conformations of CDR loops (encircled) in antibody models for failed CASP15 antibody-antigen complexes. Experimental structures are in gray, and model structures are in red and magenta
